# ninjaCap: A fully customizable and 3D printable headgear for fNIRS and EEG brain imaging

**DOI:** 10.1101/2024.05.14.594159

**Authors:** Alexander von Lühmann, Sreekanth Kura, W. Joseph O’Brien, Bernhard B. Zimmermann, Sudan Duwadi, De’Ja Rogers, Jessica E. Anderson, Parya Farzam, Cameron Snow, Anderson Chen, Meryem A. Yücel, Nathan Perkins, David A. Boas

## Abstract

**Significance:** Accurate sensor placement is vital for non-invasive brain imaging, particularly for functional near infrared spectroscopy (fNIRS) and diffuse optical tomography (DOT), which lack standardized layouts like EEG. Custom, manually prepared probe layouts on textile caps are often imprecise and labor-intensive.

**Aim:** We introduce a method for creating personalized, 3D-printed headgear, enabling accurate translation of 3D brain coordinates to 2D printable panels for custom fNIRS and EEG sensor layouts, reducing costs and manual labor.

**Approach:** Our approach uses atlas-based or subject-specific head models and a spring-relaxation algorithm for flattening 3D coordinates onto 2D panels, using 10-5 EEG coordinates for reference. This process ensures geometrical fidelity, crucial for accurate probe placement. Probe geometries and holder types are customizable and printed directly on the cap, making the approach agnostic to instrument manufacturers and probe types.

**Results:** Our ninjaCap method offers 2.2±1.5 mm probe placement accuracy. Over the last five years, we have developed and validated this approach with over 50 cap models and 500 participants. A cloud-based ninjaCap generation pipeline along with detailed instructions is now available at openfnirs.org.

**Conclusions:** The ninjaCap marks a significant advancement in creating individualized neuroimaging caps, reducing costs and labor while improving probe placement accuracy, thereby reducing variability in research.

## 1 Introduction

### 1.1 State of the Art and Motivation

Accurate placement and co-registration of probes and sensors on the head are crucial for non-invasive measurements of targeted neural activity in human neuroscience research. This process typically begins with researchers carefully identifying target brain areas, which then guide the layout, placement, and fixation of probes for data collection and analysis.

This applies to functional near-infrared spectroscopy (fNIRS)^1,2^ and electro-encephalography (EEG), but also to the emerging field of optically pumped magnetoencephalography (oMEG)^3^ and other head-worn sensors. In EEG, the 10-20 system and its derivatives (10-10 and 10-5) provide established, widely accepted standards for electrode placement^4^. By using predefined relative distances between electrode positions and anatomical landmarks, the 10-20 system accommodates variations in head circumferences, ensuring consistent, standardized measurements across individuals. Currently, fNIRS lacks its own standardized layouts and caps, and researchers frequently use the EEG 10-20 system as a workaround^5^. However, since the design of fNIRS probes is critical for optimal signal and spatial resolution, this approach has limitations:

1. The distances between 10-20 coordinates are *relative* and depend on the individual head circumference, but fNIRS measurements are highly dependent on *absolute* source-detector distances. Channel distance affects sensitivities to hemodynamic changes in brain vs superficial tissues, signal strength and quality. As opposed to EEG, scaling of probe geometries with head circumference is usually not desired in fNIRS. Even at a fixed standard 56cm head circumference, the possible channel distances between 10-5 coordinates do not cluster around the typical target distances for fNIRS optodes but are instead spread across a wide range around those (see **Figure 1**).
2. The 10-20 coordinates constrain the shape and layout of fNIRS probe geometries, which specifically can become an issue in high-density probes as used for diffuse optical tomography (DOT). High density grids or triagonal/hexagonal geometries, for instance, are often not compatible with the standard EEG coordinates.

**Figure 1:**
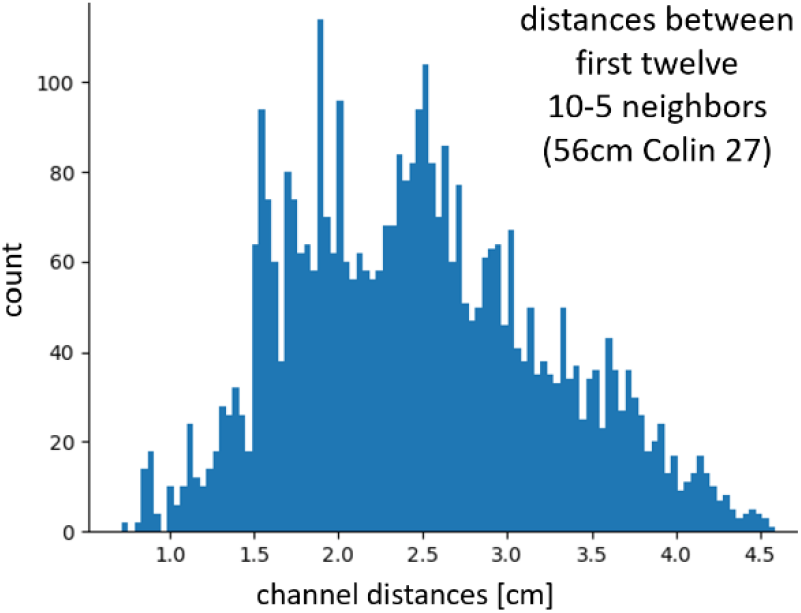
Euclidean channel distance distribution for first 12 neighbors from the 10-5 system on a 56cm Colin 27 head. Common targets for fNIRS channel distances are <1.5cm and 3-3.5cm

The common alternative is for researchers to manually prepare caps, a laborious and imprecise process that involves aligning probe layouts with (virtual) head models and making cuts in the physical textiles in the corresponding approximate positions. This approach adds variability to the exact placement of probes on the individual’s head, and further variability is added if variations of head morphology, e.g., based on ethnic backgrounds, are not sufficiently captured by the cap designs. Digitizing and co-registering sensor positions on participants’ heads during experiments, e.g. with photogrammetric approaches^6,7^, allows the assessment of how well the actual anatomical and targeted positions are matching. However, if mismatches are identified, it is usually a cumbersome and iterative manual process to improve the probe layout on the cap. Moreover, even when a subject specific head model (e.g., a structural MRI scan or a photogrammetric surface scan) and individual target regions are available, translating this information onto the physical cap is not straightforward.

To address these challenges and to leverage the advantages of widely available advanced rapid prototyping technologies, employing 3D printing for the creation of customized head caps presents a compelling solution that we have explored and refined over the last years.

### 1.2 Challenges in 3D printing head caps

What are the challenges in 3D printing head caps for neuroscientific studies? 1) Printing flat objects that accurately conform to the curvature of the head and offer a uniform fit, is a non-trivial task. Printing a whole (non-flattened) cap is technically unfeasible for most flexible materials and most users’ printers. Mapping the 3D shape of the head onto a 2D printing plane requires careful consideration of relaxation and deformation strategies and the right choice of the cap’s lattice structures. 2) Materials and manufacturing present another set of challenges: Materials that offer the necessary stretchiness and durability need to be identified. The manufacturing process itself may involve manual labor, adding complexity to the production workflow. 3) Figuring out the best way to split the head into panels and how those panels are later reunited requires consideration of the print bed size and minimizing deformation, especially when dealing with three or four panels. 4) In the case of high-density (HD) and ultra-high-density (UHD) fNIRS caps, general challenges are to 4.1) obtain uniform coverage, minimizing gaps between tiles or patches, to 4.2) optimize sensitivity, uniformity, resolution and minimize localization errors on the brain, while 4.3) meeting geometrical constraints, e.g., feasible channel distances that take into consideration both obtainable signal quality and mechanical constraints from optodes and holder sizes.

### 1.3 Our Solution

Solving the above challenges and incorporating many years of our experience in head-gear and probe designs^8,9^, we have created a method and process that enables researchers to fabricate 3D printed caps with fully customizable integrated holders, eliminating the need for laborious manual cutting and preparation of standard EEG caps (see **Figure 2**). We call these “**ninjaCaps**”.

**Figure 2:**
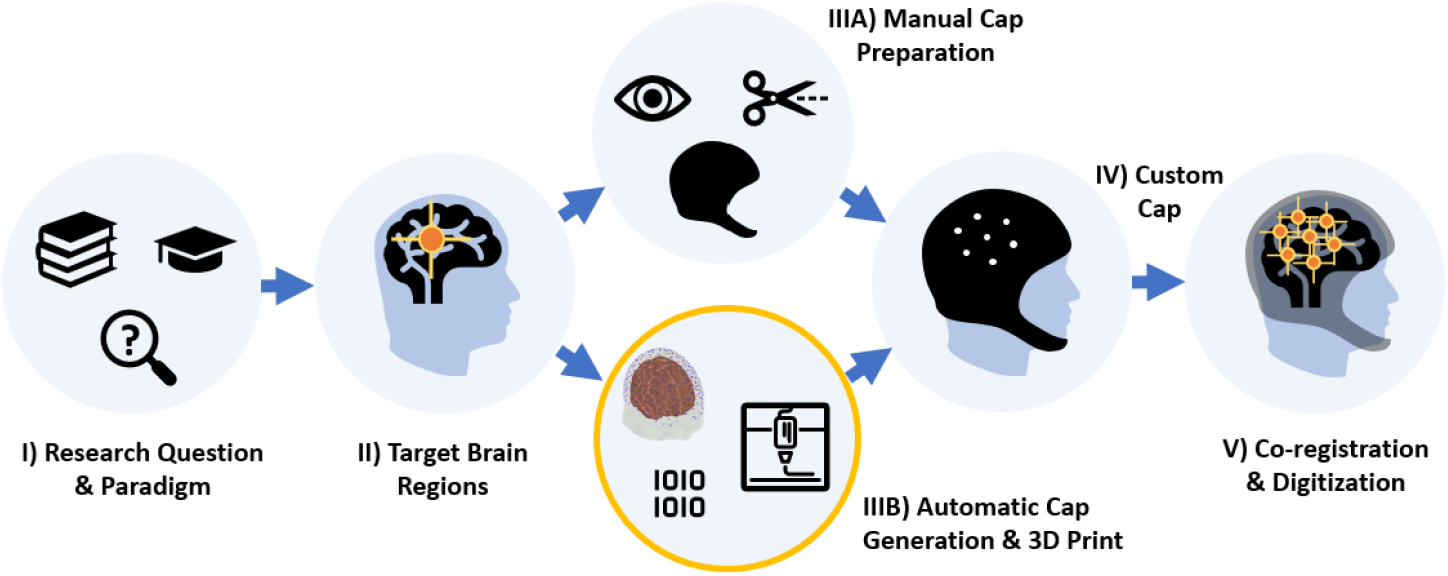
Conventional process from paradigm to registered measured brain activation. New step IIIB): Automatic generation and 3D printing of caps for customized probe layouts and head anatomies, instead of manual preparation of standard textile caps (IIIA).

This approach can significantly reduce both the time and cost associated with cap preparation, making it more accessible and affordable for researchers to conduct customized studies. Each experiment and even individual can have a dedicated cap, eliminating the need for reusing and re-assembling caps, thereby enhancing experimental control, and reducing potential sources of variability. The reproducibility of experiments can be improved, establishing a “virtual ground truth” through the standardized production of 3D printed caps. The versatility of 3D printing enables the creation of customized and complex cap designs that can integrate any type of head mounted sensor, for instance fNIRS optodes, EEG electrodes, or other sensors or components such as accelerometers, in almost any geometry and mixture. This integration enhances mechanical stability and expands the range of potential applications in research settings. Lastly, researchers can tailor the cap design to fit individual head sizes and shapes accurately, improving the quality and reliability of measurements. Rather than relying on standardized head models, the individual head geometry can be precisely replicated, ensuring optimal contact between sensors and the scalp.

In this manuscript, we detail our approach step-by-step, elaborate on innovations and lessons learned, and share resources to enable the community to create their own 3D printed caps.

## 2 Methods

### 2.1 Overview of ninjaCap Generation

We use our established *MATLAB*-based brain atlas software AtlasViewer^9^ for probe layout design and for mapping of sensor positions to the brain surface. Also in *MATLAB*, we project from curved head surfaces to 2D panels with a spring relaxation algorithm and use hex lattice structures to obtain better deformation and uniform fit. We then automatically extrude and assemble panels and holders in the open *Blender3D* (Blender Foundation) software and 3D print the caps using flexible thermoplastic polyurethan (TPU) filaments. Our material of choice in this work is *ninjaFlex* (Fenner Precision Polymers). The cap panels are then assembled with an ultrasonic welder. An overview of this process, which will be described in detail in the following sections, is shown in **Figure 3**.

**Figure 3:**
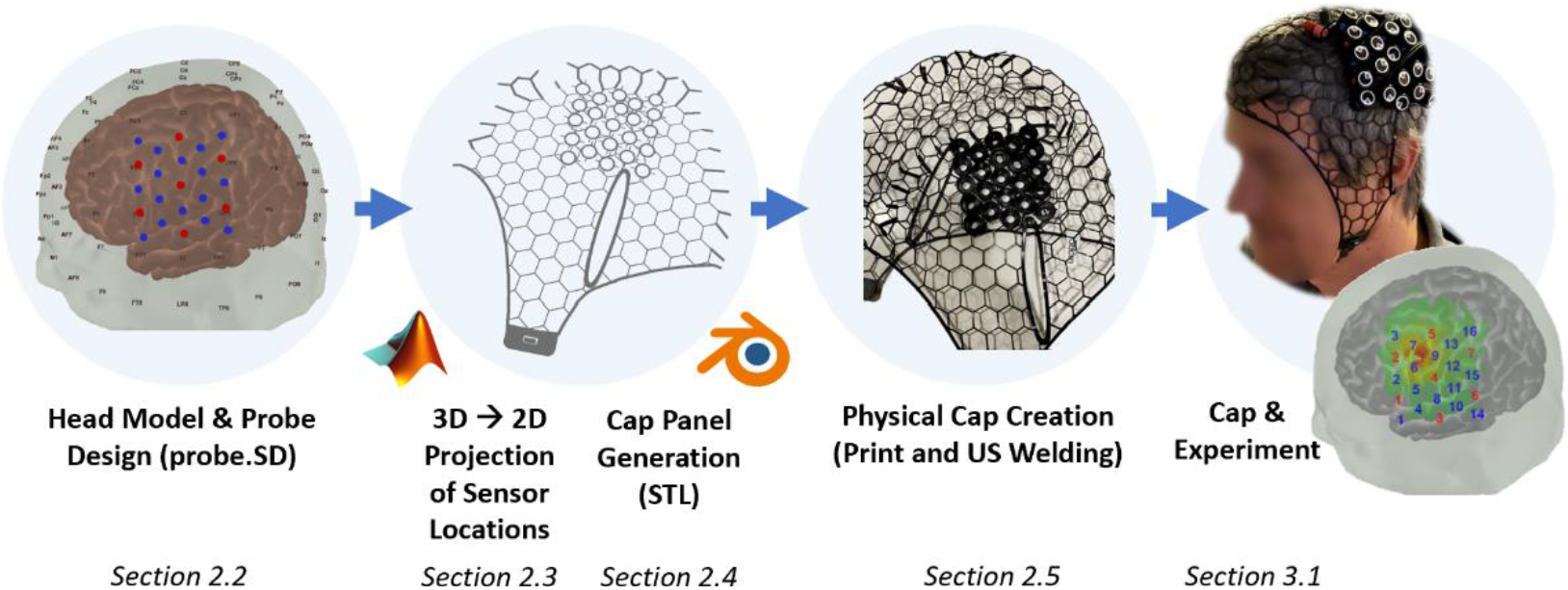
Visualization of ninjaCap generation from probe design to output

To make the ninjaCaps available to the community, we are hosting an automated pipeline for cap generation on bfnirs.openfnirs.org. The input to this cloud-based pipeline is the desired head circumference (HC) of the cap together with a “Probe.SD” file that can be generated in AtlasViewer. The output is a set of four .STL files (cap panels) that can be 3D printed with flexible TPU and assembled by the user.

### 2.2 Head Model & Probe Design

ninjaCap generation is based on two key input elements: A 3D head model and a probe design specification. The head model provides anatomical guidance during probe design to target the desired brain areas, and the head surface geometry is used for the generation of the cap panels. The probe design file includes the geometrical constraints (e.g., for anchoring), arrangement (e.g., source-detector separations) and types of probes (i.e. specifying the corresponding holders/grommets) that are then registered onto the surface of the head model. **Figure 4** visualizes source-detector arrangement, necessary anchoring information for probe registration, and the type of the grommets (e.g. fNIRS optode or EEG electrode).

**Figure 4:**
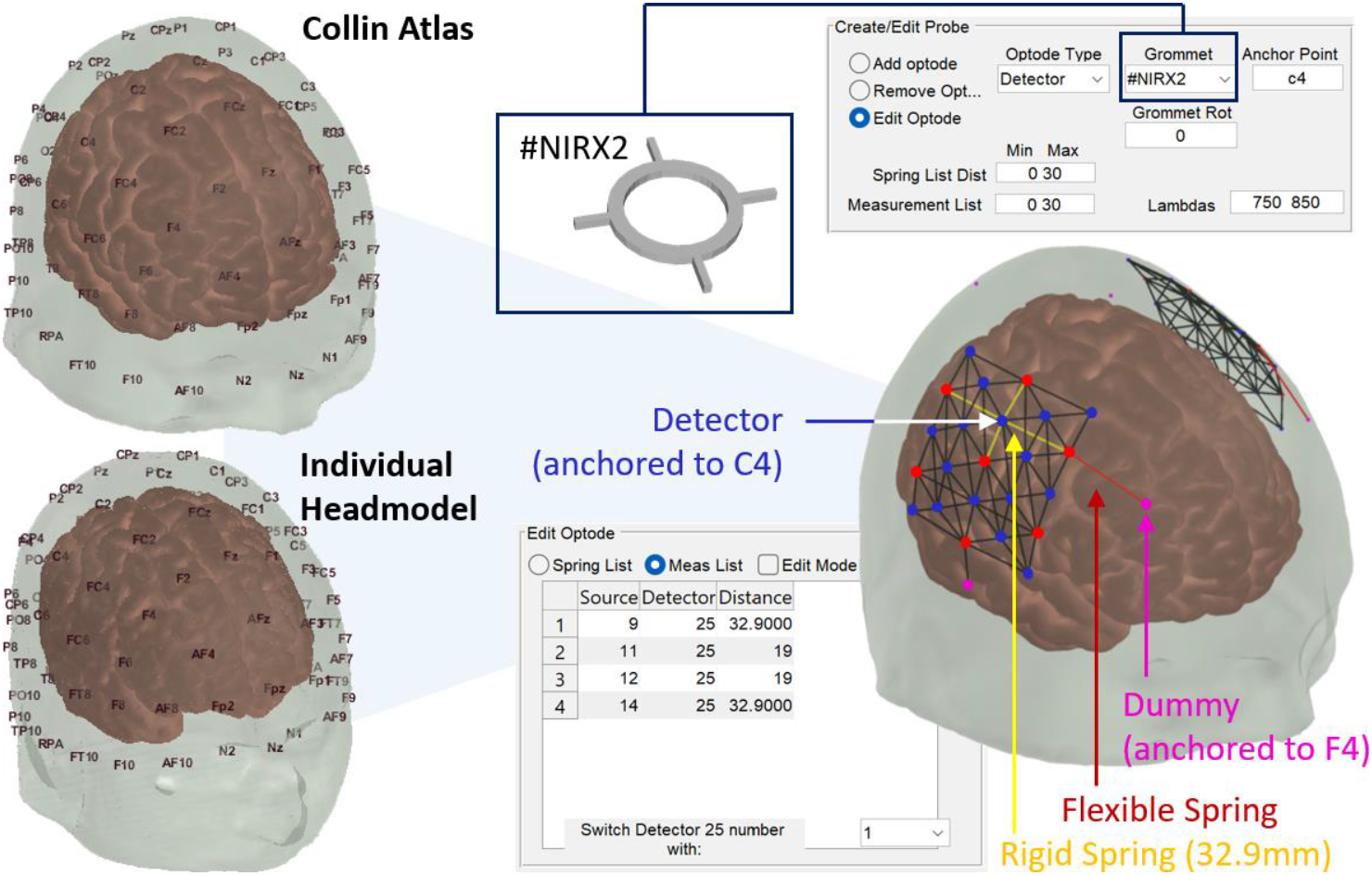
Head-model based probe generation and registration in AtlasViewer. Bilateral HD-DOT patches, each with 7 sources and 16 detectors that are inter-connected with rigid springs. Patches are flexibly connected to anchor points. In this example, optodes have grommet types “#NIRX2” to allow for the placement of NIRx optodes on the cap. Windows show configuration details for the selected detector on C4.

#### Head model

Our default surface model in AtlasViewer is the widely used Colin 27 head model^10^. Other atlases or individual MRI anatomies (e.g., NIFTI format) can be used as well, see ^*,9^ for a detailed tutorial for how to import them. The current ninjaCaps pipeline on bfnirs.openfnirs.org does not yet natively support user-defined head models, but the cap generation pipeline and following process is agnostic to the head model used. Before cap generation, the head circumference (HC) has to be specified. We currently scale the head model linearly only with HC, but other distances can also be employed (in AtlasViewer: Iz to Nz distance and RPA to LPA distance).

#### Probe design file

Users can design their probes in AtlasViewer, which are then saved in a “.SD” MATLAB file. Key elements of the .SD file for cap generation are a list of 3D node positions, anchor points, springs, 10-5 landmarks, and the head mesh to be used. Since AtlasViewer is designed for fNIRS, nodes are called optodes in the interface, but can represent any other type of sensor as well. Available node/optode types are “source”, “detector” or “dummy”. Source and detector type assignments are relevant to constructing an fNIRS probe and channel measurement list that is relevant for acquisition and plotting of fNIRS signals, but not for the generation of the physical cap itself. All three node types always have a “Grommet Type” assigned to them. Grommet types (in the format “#<abcde>) identify the fully customizable 3D holders that will be printed on the cap. Nodes/optodes are interconnected by “springs” that are either flexible and can be stretched, or rigid and have a fixed length which constitutes a geometrical channel distance constraint. This spring-based approach enables projection and registration of the probe geometry on the head surface. Another constraint during probe registration is generated by anchors: Nodes can be anchored to 10-5 landmarks on the head, which enforces the node position to remain exactly on this landmark during spring-relaxation. The user typically assigns such anchor points to a small set of nodes/optodes that are to be fixed to their corresponding anatomical landmark. When scaling the head model, the subset of grommets/dummies that are anchored to these relative locations scale with the head. In contrast, any distance between optodes constrained by a rigid spring is a fixed parameter that does not get scaled but remains constant to maintain the specified probe geometry. Two examples: 1) A cap can be generated with only “dummy” nodes that have EEG-electrode grommet types assigned to them. Anchoring these to standard 10-20 positions will result in a 3D printed standard 10-20 EEG cap. 2) To place a HD-DOT geometry with fixed inter-optode distances over the corresponding somatosensory area for a finger tapping experiment, one could anchor one measurement optode to the C3/C4 10-20 position, and at least three “dummy” optodes with springs without fixed distance constraints to 10-20 landmarks to further constrain the HD-DOT patch placement. **Figure 4** shows an example for such a bilateral HD-DOT probe registered to the Colin atlas and provides details for its configuration. Please see ^9^ and ^†^ for a detailed tutorial on how to design 3D source detector geometries and the corresponding SD file in AtlasViewer.

### 2.3 Flattening and Projection of Sensor Locations

With the curved head model with co-registered probe positions available, the crucial next step is to create flat panels from this head model that can be 3D printed. The panels need to yield an accurate placement of each individual sensor holder on its intended position on the curved surface of the head after cap assembly. In our first ninjaCap versions, our approach was to calibrate points between the curved head and a flat template to then stretch flat panels to fit over the curved head surface. However, iterating over several cap generations has led to a different and more precise approach to project from 3D positions to 2D cap panels, depicted in **Figure 5**.

**Figure 5:**
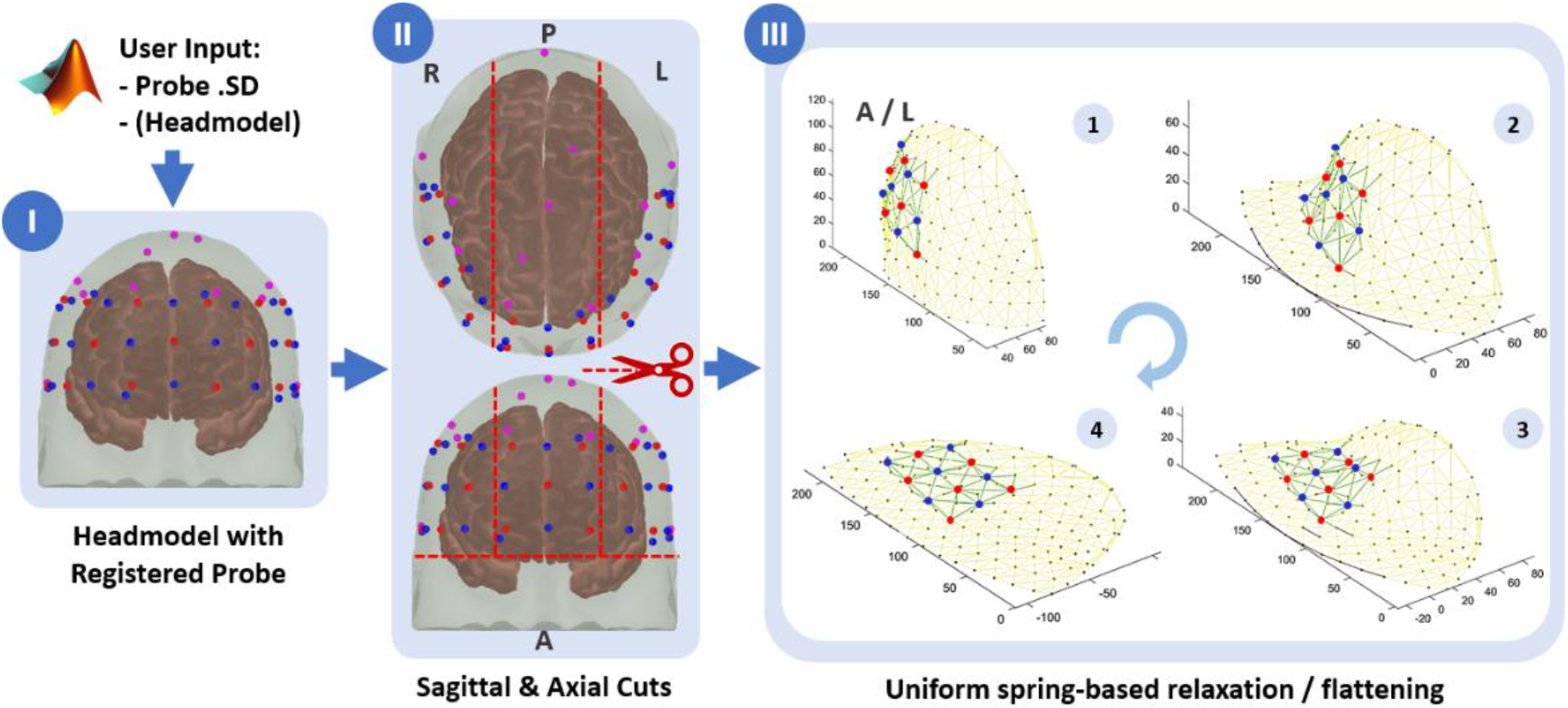
**I)** head model with a co-registered pre-frontal fNIRS probe. **II)** Cutting into panels. **III)** Flattening panels with iterative spring-based relaxation (steps 1-4). Flattening is shown for the panel of the left (L) side of the head.

A prerequisite of the following approach is that all the 10-5 points as well as the sensor/dummy nodes from the registered probe on the head are first incorporated as vertices into the surface mesh of the head.

First, the 3D head mesh is cut into 4 pieces (see **Figure 5 II**): An axial cut through Inion (Iz) and 15mm above Nasion (Nz) that discards any part of the head mesh below the eyebrows (including neck). With the head centered at the Iz-Cz-Nz midline, two sagittal cuts are made on either side so that the middle part has a width of 3/7th of the total distance between LPA and RPA through Cz. These sagittal cuts go very close to the reference points NFP1/2, NFP1/2h, I1/2 and I1/2h of the EEG 10-5 system and cut the upper part of the head into three parts (left side, mid, right side) that are then used for the generation of the left, right and mid/top panels of the cap.

The resulting three surfaces are then individually flattened and projected to 2D (see **Figure 5 III**) following an iterative procedure that treats the edges between vertices of the 3D meshes as springs. During this procedure, distances of sensor/optode node vertices relative to their closest three 10-5 vertices are kept fixed: 1) We apply a normalized gravity force along the Z axis that pulls all vertices towards the X-Y plane at Z=0 with a very small displacement, normalized to a maximum of a 0.1mm displacement per step. 2) We then apply spring forces that act on the relative distances in the curved mesh as consistently as possible, leading to a **uniform distortion**. 3) We repeat steps 1) and 2) until all vertices’ Z coordinates are approximately at 0. Holding the total area of the surface nominally constant when flattening ensures that positions on the flat panels end up in the corresponding curved positions on the head when all panels are re-assembled in a 3D cap. This approach preserves the local distances between 10-5 reference points, which also locally preserves area and thus makes the area of the printable flattened panel and the area of its corresponding 3D head piece roughly identical. Since optode/electrode positions are vertices in those meshes, the flattening transformation also yields the correct locations of the corresponding grommets to be placed on the 2D cap panels. **Box 1** shows the pseudocode for the key steps of the transformation. The supplements to this article provide two videos that visualize the flattening and spring-relaxation process for a side and a mid panel.

#### Box 1

**Pseudocode for spring-relaxation approach to obtain flattened 2D panels**

`mapPanelandGrommets()` prepares the 3D mesh for flattening by setting up the vertices and edges for spring relaxation and identifying key reference points.

- **Inputs**: list and 3D positions of 1) vertices and edges of the cut mesh piece belonging to the panel to be generated, 2) grommet positions and 3) 10-5 reference points.
- **Outputs**: 2D positions of the mapped vertices for panel, grommets, and reference points.
  - **Initialization**: Prepare 3D vertices for spring relaxation:
    - Calculate equidistant points along the panel’s outline.
    - Identify relevant reference points and labels.
    - Create list of vertices (`vHex`) and edges (`eHex`) for spring relaxation.
  - **Create New Vertices and Edges (Connections)** for
    - reference points
    - outline seam points, and for seam with reference points
    - grommets and their closest two reference points and add them to the list of existing vertices and edges.
  - **Run and Repeat Spring Relaxation**: Invoke `SpringRelax_func` iteratively until all vertices’ coordinates are on a 2D plane with Z ≈ 0.

`SpringRelax_func()` performs the spring relaxation algorithm, adjusting the 3D vertices to flatten the mesh onto a 2D plane.

- **Inputs**: Vertices (`vHex`), edges (`eHex`), and desired edge lengths (`hHex`).
- **Output**: Adjusted vertices (`vHex`), edges (`eHex`) after spring relaxation step.
  - **Spring Model Setup**: Calculate initial Euclidean lengths of the edges in the mesh.
  - **Apply Gravity Force**:
    - Apply a normalized gravity force to all vertices along the Z axis
  - **Calculate Resulting Spring Forces**:
    - For each edge, calculate the force exerted on the vertices based on Hooke’s law:
      - Calculate the difference between the current and the desired lengths of each edge.
      - Compute the forces in the X, Y, and Z directions.
    - **Update Positions**:
      - Apply a global scaling factor to the force to control the update magnitude.
      - Adjust each vertex’s position in the X, Y, and Z directions in response to its applied force.

### 2.4 Panel STL Generation and Cap Assembly

To generate and finalize printable cap panels from the flattened 2D projections several additional steps are necessary (**Figure 6**). These are: IV) Completing panel outlines. V) Filling panels with a hexagonal lattice that grommets can be attached to, and that provide structure and uniform stretch. VI) Creating panel seams and “welding tabs” for assembly after printing. VII) Extruding the 2D panel to a printable 3D mesh, adding ear slits, grommets and chin/neck strap holders according to coordinates and grommet types provided. Steps IV-VI are part of our MATLAB script; for step VII, our script makes use of the advanced 3D meshing and manipulation functionality of Blender 3D, via its internal python interface.

**Figure 6:**
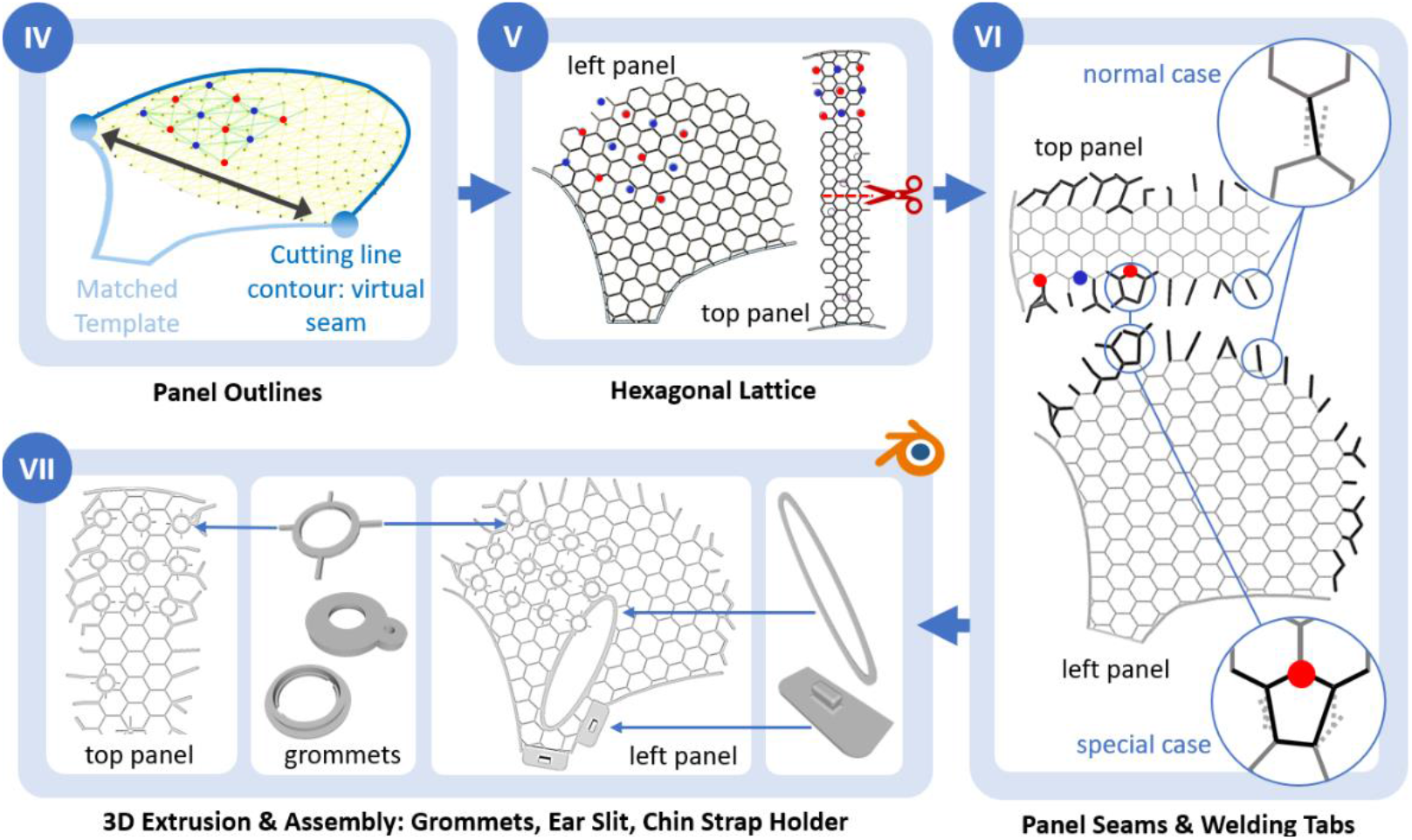
**IV)** Defining virtual panel outlines and matching a template for lower outline to side panels. **V)** Filling the panel outlines with a hexagonal lattice. **VI)** Generating welding tabs along virtual seams, considering normal and special cases with grommets close to seam. **VII)** 2D to 3D extrusion and assembly of panels in Blender3D: Adding grommet types, ear slit and chin (and neck) strap holders.

#### Panel Outlines

After flattening the meshes, we use the contours of the cut lines to generate the panel outlines. This maintains the absolute coordinates of projected grommets that will be placed later in the process. For the middle panel, the outline is defined by the sagittal and axial cutting lines after flattening. We cut the middle panel in half to enable printing on a print-bed of 30cm edge length. For the side panels, the sagittal cutting line after flattening defines the upper outline but the lower part needs to be added: it defines the cap outline underneath the chin, ear and above the neck. We use a template that we optimized over the years, it is laterally stretched to connect to the outmost points of the upper outline, see **Figure 6 IV**.

#### Hexagonal Lattice Structure

The outlines are then automatically filled with a hexagonal lattice with edge length 10mm (see **Figure 6 V**). We have previously explored different geometries such as a square lattice or zig-zag segments. Those proved less suitable to achieve uniform stretch/deformation of the cap on the head. The 10mm edge length of the hexagonal lattice is a tradeoff between good deformation (smaller hexagons reduce stretchiness of the cap) and the minimum area coverage needed to ensure that grommets are always well-connected to the lattice and do not end up in empty spaces.

#### Panel Seams

To form the final cap the printed panels will be assembled via ultrasonic welding. For this, each panel has “welding tabs” that must be aligned across the joint seam of a panel and its corresponding neighbor. Welding tabs are generated with a thicker outline that facilitates better connection during ultrasonic welding. We generate these as follows: Using the sagittal cutting line as a virtual seam between both panels, we identify all edges within a fixed physical distance directed towards the seam. For each edge we find its closest corresponding edge from the other panel and reposition both towards their shared intermediate point. Each edge of the pair is then extended out up to the end-node of the other. Both together constitute the overlapping “welding tabs” for manual assembly (see next section). Two special cases need to be considered. 1) If one panel has more intersecting edges towards the seam than the other, we redirect and join two “welding tabs” from one panel to a single corresponding one from the other. 2) Grommets can be close to or overlapping with virtual seams. If a grommet’s location is within one hexagonal edge length of the virtual seam, we extend the overlapping edge to the next node on one panel side. Please see **Figure 6 VI** for visualizations.

#### 2D to 3D Extrusion and Assembly of Ear Slits, Grommets and Holders

The resulting two side panels and two halves of the top panel, all with their generated outlines, hexagonal lattices and welding tabs are saved as 2D .STL files. Together with the corresponding grommet types, 2D coordinates and rotation in a .digtpts textfile these are processed by a Blender 3D python script to assemble the final panels for printing (see **Figure 6 VII**). We use Blender’s own Python with its native functions for manipulation of meshes, e.g. boolean joining and cutting out of objects and some conventional modules (e.g., SciPy) installed. The 2D panels are first extruded by 0.9mm to create printable 3D volumes. For each listed grommet type, its corresponding STL file is loaded from a library folder and placed and merged to the panel’s hex lattice at its corresponding 2D coordinate. Using the 4-letter identifier #<abcde> in the probe .SD design file as a naming convention (STLs are always named “grommet.stl” saved in a folder with the same name as the identifier) enables easy adaptation and placement of fully customized 3D holders and objects on the panel. Standard chin and neck-strap holders are placed at pre-defined positions on the side-panel. Finally, ear slit holes are stamped out of the side panels and corresponding outline templates are placed. The ear slit outlines are centered at 15mm up from LPA and RPA 10-20 reference points. These relative positions are the result of ongoing validation and user feedback to optimize fit and comfort. The whole process generates four cap panels: The left- and right-side panel, and one mid panel that is cut in half to make it fit on print beds with 30cm edge length.

### 2.5 Physical Cap Creation

The four resulting cap panel STL files are now ready to be printed with any 3D printer with a print bed >30x30cm^2^ that supports printing with thermoplastic polyurethane (TPU) flexible filaments such as ninjaFlex. Printing thin features using this filament with the quality needed for our head caps depends heavily on a few settings. These are primarily the extruder temperature, nozzle offset, and the line-to-line overlap. The temperature of the extrusion must be tailored to the thermal transfer properties of the nozzle on the printer. The highest quality combination for printing we found is to use a hardened steel nozzle at 225 degrees Celsius and a print speed of 10-12 mm/s. Using nozzles made of either plated brass or copper requires the temperature approximately 10-15 degrees lower or the print speed to be increased a proportional amount. If the extrusion temperature is set too high, the final part will have voids in it due to the filament leaking out of the nozzle as it travels, weakening the part. This can be identified by strings of the TPU crossing areas of the part that should not have any material. The nozzle offset needs to be dialed in for each printer and should be set close enough that the deposited line is compressed into the build plate such that the resulting line is about 1.5-2 times as wide as it is tall. This setting combined with a line-to-line overlap of 20% ensures that the individual lines on each layer are combined enough for the top and bottom layers to no longer show distinct paths of travel on a finished print. If the overlap is insufficient, the finished cap will experience weak adhesion and wear out prematurely.

The printed panels are then manually assembled into a cap. This is done by using cost-efficient low-power handheld ultrasound (US) welding devices such as the one depicted in **Figure 7 IX**. We use a QUPPA clamp device, which is a 20W 57kHz countertop ultrasonic sealer to make welding points on PET, PS, PVC, and PE materials. To connect panels along the seam, the corresponding welding tab pair of each panel is placed on top of each other and then welded by gently pressing down with the US welding clamp. Welding time in seconds (set by the US welding device) and manual pressure at the welding clamp determine the quality and durability of the welding points along the seam. If pressure and/or welding time are too low, the TPU does not connect properly; if too high, the welding tabs can get destroyed. With the right settings, the resulting connections compare with properties (stretch, durability) of any other part of the 3D printed TPU mesh. After initial training, assembling a cap takes approximately 0.5-1 hr. As this cap already has all grommets printed at the desired locations, it is ready for population and experimental use right away.

**Figure 7:**
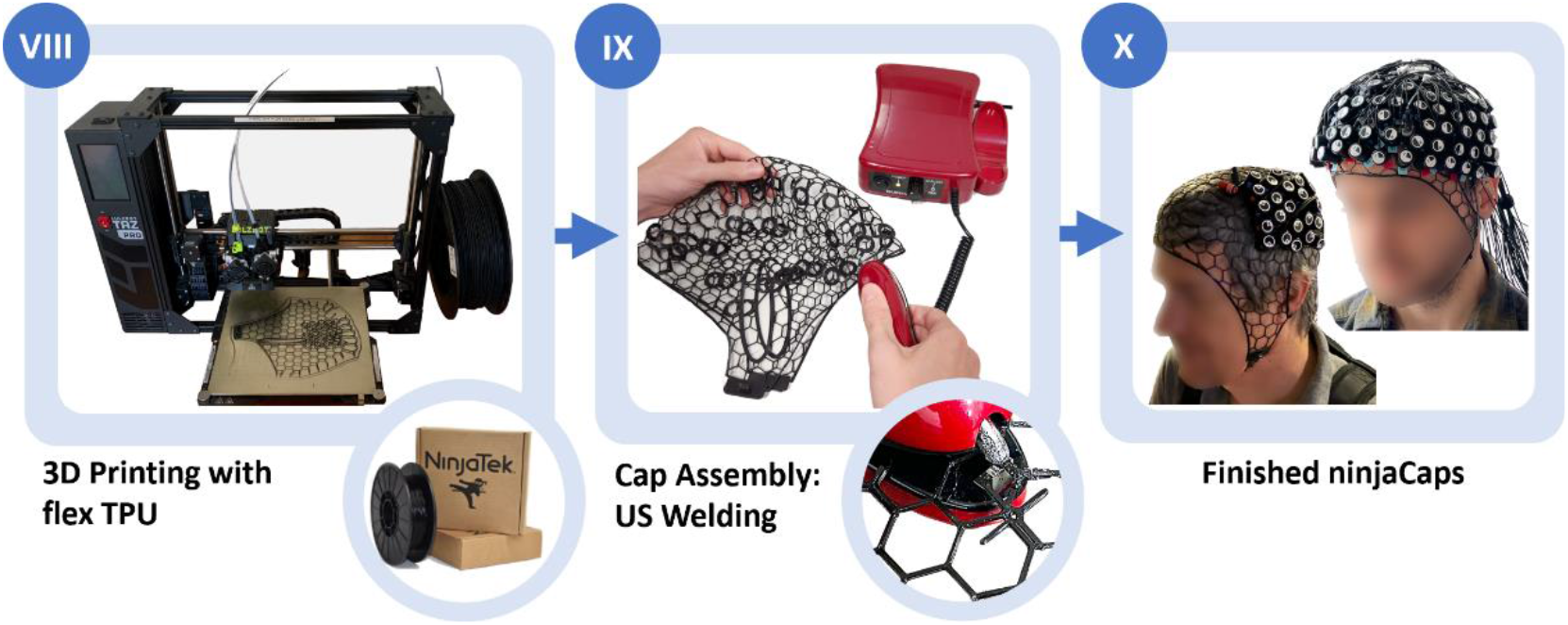
**VIII)** 3D printing of 4 cap panels. **IX)** Manual assembly using ultrasonic welding clamp. **X)** Finished ninjaCaps.

### 2.6 Verification and Validation

The presented methodology for creating ninjaCaps has been refined over the last five years through an ongoing process of development, verification, and validation with several hundred participants. This process evaluated the accuracy of probe placement, along with the comfort, fit, and longevity of the caps (see next section). To quantify the precision of our technique — translating 3D target coordinates on a head model to flat, printable cap panels, and subsequently to the physical probes at the target coordinates on the actual head — we employ the following procedure: Utilizing the Colin 27 56cm head circumference head model with known EEG 10-20 coordinates, which serves as our ground truth, we (1) fabricate ninjaCaps with grommets positioned at these coordinates, and (2) create and print a version of the Colin head model that features small indentations that denote the correct positions. Using the cap on the head we then measure and report the distances between the centers of each grommet on the cap and the corresponding indentation on the model across the 10-20 positions.

## 3 Results

### 3.1 The ninjaCap

Over time, starting in 2018 we have evaluated 20 models in the first generation (template based panel generation), and then 50 models since August 2022 in the second generation that follows the spring-relaxation-based approach in this manuscript. **Figure 8 A)** shows three such cap variants for fNIRS and HD-DOT. Mixed EEG-fNIRS caps and EEG-only caps have also been constructed. Because of the limited size of an average 3D printer’s print bed, side panels and the split mid panel need to be printed sequentially in three stages. Each print takes between 1hr 45min and 2 hrs 30min for the side panels (dependent on cap size) and between 1hr 15min and 2hr for both mid panel pieces combined. Currently, the overall cost of creating a cap is ∼5-7 hrs total printing time, 30 cm^3^ ninjaFlex volume (ca. $6.00) and 1.5-2 hrs of manual labor to print, clean and assemble the panels. The current versions of our caps have been successfully used an average of cycles before wearing out or breaking.

**Figure 8:**
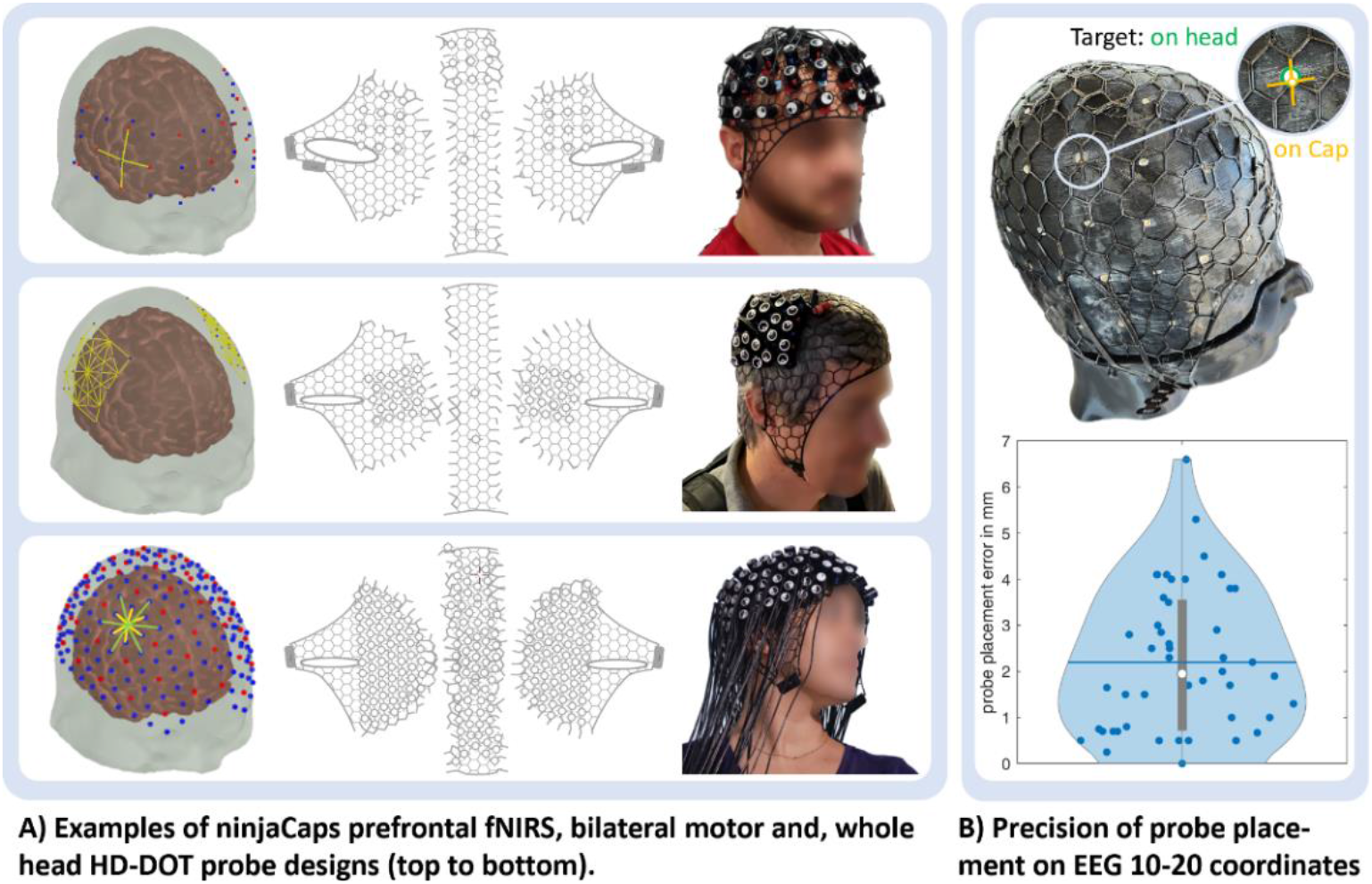
**A)** Examples of three fNIRS probe types and ninjaCaps. First row with improved ear slit design. **B)** Violin plot of ninjaCap probe placement errors on 3D printed ground truth head model with EEG 10-20 positions. Blue dots: individual placement errors. Blue line / white dot: mean / median error. Grey vertical bar: Interquartile range

### 3.2 Verification and Validation

#### Probe placement precision

When evaluated on the 56cm head circumference 3D printed ground truth head model with 10-20 marker indents, our ninjaCap approach yields an average probe placement error and standard deviation of 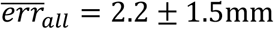. **Figure 8 B)** shows the corresponding violin plot. The error is not evenly distributed across the whole head surface, but is lower towards the medial and anterior regions of Fz, Cz (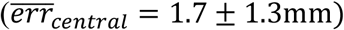), constituting most points at the lower end of the distribution, and larger towards the lateral and dorsal regions of F7/8, P7/8 and O1/2 (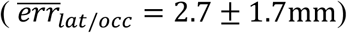), constituting most points at the upper end of the distribution.

#### User testing – fit and comfort

To date, more than 50 ninjaCaps have been tested by more than 10 independent research groups with 500+ participants. At early stages of the cap design, some users reported discomfort caused by the caps’ ear slits, particularly for longer studies. This feedback was incorporated by testing different shapes, positions, and angles with our user base, resulting in a widened and backwards-slanted ear slit with the position and angle reported in this manuscript. **Figure 8 A)** shows both old and latest ear slit solutions (1^st^ row, prefrontal probe: latest ear slit; 2^nd^ and 3^rd^ row, bilateral and whole head probe: older ear slit). Additional user feedback led to the addition of an optional strap similar to a chin strap that runs to the back of the head that can improve fit and uniform stretch on the head. The quality and amount of feedback received since then indicates comfort and performance comparable to other solutions (such as Easycaps) that our users are accustomed to.

#### Durability and mechanical lessons learned

To improve durability of the caps, we iterated on the thickness (number of layers) of the lattice of the printed panels. Too thin prints with lattice thickness ≤ 0.7mm broke too easily and stretched too much after a few (∼10) uses; too thick prints did not conform well to the head. The best tradeoff between both was found to be a thickness of 0.8mm, which has been used ever since. Similarly, to optimize durability and stretchiness, we ended up using lattice (=hexagon edge) lengths of 10mm and lattice widths of 0.85mm. For robust connection of panels using the ultrasonic welder, we ended up with an optimal welding tab width of 2mm. Another crucial set of parameters that was optimized is the 3D printing profile, that depends on material and printer used. We provide the 3D printing profile for LulzBot TAZ Workhorse printers in the forum on openfnirs.org on request. Based on our and our user base’s experience with caps manufactured with these parameters, the number of usage cycles/lifetime of ninjaCaps is estimated to be comparable to conventional textile caps such as Easycaps.

### 3.3 Availability

To make the ninjaCaps available to the community, we are hosting an automated pipeline for cap generation on bfnirs.openfnirs.org. The pipeline is based on the process described above and requires a .SD probe specification file. Generated printable cap panels can afterwards be downloaded as .STL files. Probes can be generated with AtlasViewer, which is openly available on github.com/bunpc/atlasviewer as a MATLAB repository together with comprehensive tutorials. Some ninjaCaps variants for fNIRS and HD-DOT that we generated over the last years are also openly available for download on openfnirs.org/hardware/ninjacap.

## 4 Discussion & Conclusion

### 4.1 Alternative Approaches

On our path to the current ninjaCap generation we considered alternative approaches to generate custom fNIRS/DOT headgear and caps. A brief summary of alternatives follows to contextualize and motivate the solution that we presented in this manuscript.

#### Combining 3D printed probe holders with textile caps or custom headgear

Custom holders or modules can be 3D printed and then manually incorporated into textile caps,^11,12^ or can be used to constitute mechanical elements of headgear.^8,13,14^ However, this approach is labor intensive not readily scalable.

#### Direct 3D Modeling and Printing

Caps can directly be designed in a 3D environment, considering the curvature and specific anatomical features, and then 3D printed in a rigid or even flexible material^15^. As in our approach, individual surface anatomies, e.g., from a structured MRI or a photogrammetric scan, can be used to design subject specific caps or helmets – however, without the need for projecting onto a 2D plane. This has the potential of providing even higher precision, especially for rigid structures and helmets that can support heavy probes. To print flexible 3D caps however, special 3D printers are needed, because extruded TPU/ninjaFlex has non-isotropic mechanical properties. The strength in the “z” dimension between layers is not as high as within a layer, especially in the direction of the strands and compromises in flexibility/stretchability of the material must often be made.

#### Template-Based Panel Generation

Instead of projecting the 3D surface to 2D, one can use pre-defined templates of head shapes and sizes to generate 2D panels with a set of positions (e.g. 10-5 landmarks) calibrated to the 3D surface. While easy to implement, a drawback is the reduced accuracy of probe positions, especially when individual head geometries are to be used. In our ninjaCaps, we started with this template-based panel generation and progressed towards the presented spring-relaxation algorithm presented in this manuscript to improve precision.

#### Conformal Mapping

Projecting the 3D head surface onto a 2D plane with minimal distortion can be accomplished with the technique of conformal mapping, known from cartography. Conformal mapping is excellent for locally preserving angles and shapes at small scales but does not necessarily preserve the overall lengths or areas. While a powerful mathematical tool, it is therefore not the most suitable approach for this specific application, as preserving exact lengths (channel distances) is a priority.

### 4.2 Further Improvements and Outlook

We are continuing to expand and improve our approach beyond the current ninjaCap generation. One in-progress work is the design of an expandable cap that can be size adjusted along the sagittal midline (one size fits all), which is a requested feature based on user feedback. This updated design would also speed up the total printing time to under 6 hours and will make the same cap more adaptable to fit different subjects. Adjusting the cap size is particularly useful in High-Density Diffuse Optical Tomography experiments, in which whole-head designs (such as the one depicted in **Figure 7**) contain up to several hundred optodes and swapping optodes across caps with different head sizes can be time consuming. On the other side of the spectrum, to further facilitate the creation of subject-specific caps, photogrammetry can be incorporated to quickly create individual head scans that can then be used in the ninjaCap pipeline.

The verification of probe placement precision using a 3D printed ground truth head model showed that placement errors are small but can be further improved especially for lateral / dorsal regions of the head. There are two main sources of error in these regions. 1) Using a chinstrap with any cap, be it textile or TPU-based, leads to a localized non-uniform stretch that has a higher effect on these regions. This error could be reduced by factoring in the stretch in the probe placement before printing. However, this solution comes with the cost of an increased error whenever a chinstrap is not used. 2) We also noticed that our spring-relaxation-based panel flattening approach can lead to deformations that can contribute to a greater error in probe placement towards the lower panel seams. We are currently working on improving the flattening approach to further reduce any negative impacts and will continue to evaluate performance across different head circumferences.

Lastly, we notice that a head model that is more suitable to fit many user’s head shapes can likely further improve performance. By default, we currently generate caps based on the widely used Colin27 head model, which is not equally suitable for all subjects, as it is based on only one individual head anatomy. Head shapes differ across individuals and ethnicities, most visibly on the medial axis and particularly towards the back of the head. If no individual head model is available, a more suitable general head model directly also improves cap performance.

Introducing 3D printing of flexible head caps to the field of brain imaging offers exciting possibilities for customized and tailored data collection. This approach enables the creation of head caps that perfectly match the individual’s head shape, optimize scalp contact, ensure precise optode/electrode placement, and thus enhance the signal quality, accuracy and reliability of data acquisition while also improving wearing comfort. It streamlines the cap preparation process, enhancing the efficiency and consistency of data collection both in group studies across diverse populations and in personalized brain imaging, and therefore reduces variance and facilitates reproducibility in neuroscience research. Utilizing 3D printing technology, researchers are not only able to design and produce head caps with precise electrode/sensor placement but are also enabled to integrate multiple sensor types in the same cap for novel multimodal brain imaging approaches.

By sharing this tutorial and online resources for the ninjaCaps, we hope to aid the community to adopt these promising advances for their neuroscience research. We encourage interested readers and users to use the forum on openfnirs.org to ask for any details that they might find useful or missing.

## Supporting information

SUPPLEMENT1_Section2.3_capMappingtop.mp4

SUPPLEMENT1_Section2.3_movie_capMappingsideLeft.mp4

## Acknowledgments

The development of ninjaCap has been funded in part under NIH U01EB0239856. AvL gratefully acknowledges funding from the German Federal Ministry of Education and Research under the grant BIFOLD23B.

## Footnotes

* https://github.com/BUNPC/AtlasViewer/wiki/Importing-Subject-Specific-Anatomy

† https://github.com/BUNPC/AtlasViewer/wiki/Designing-3D-Source-Detector-Geometry-in-AtlasViewer

